# Inter-variety competition dynamics in US inbred and hybrid maize

**DOI:** 10.64898/2026.02.26.708322

**Authors:** Aimee J Schulz, Martin O Bohn, Peter Bradbury, Dayane Cristina Lima, Natalia De Leon, Sherry Flint-Garcia, James B Holland, Nicholas Lepak, Aaron J Lorenz, M Cinta Romay, Candice N Hirsch, Edward S Buckler, Kelly R Robbins

## Abstract

Variety mixtures provide a potential avenue in US cropping systems to improve yield stability and disease resistance. However, implementation of variety mixtures requires an understanding of the competitive dynamics of the crop. In this study, we examine the effects of plant competition both between and within plots through five unique experiments: 1) 5,000 diverse inbred lines in single-row plots, 2) hybrids in two-row plots developed from the above inbred lines, 3) over 4,000 hybrids measured in 141 locations in two-row plots as part of Genomes to Fields, 4) mixtures of two hybrids within a two-row plot planted across two years and five locations, and 5) mixtures of up to twenty hybrids in four-row plots in three locations. Across all experiments, we find that competitive interactions are extremely limited. Within inbred lines, height of the neighboring plot accounts for 1.2% of the variance in focal plot height. Similarly, neighbor height explains 1.7% of the variance in focal plot yield in hybrids developed from the inbred lines. The genetics of neighboring plots explains 1.55% of the variation in yield across 141 location-year environments, reinforcing the generally modest impacts of neighbor competition. In evaluating mixtures of hybrids in both two and four-row plots, we observe no yield penalty compared to conventional single hybrid plots, even with large height differentials of the hybrids included in the mixture or in mixtures of up to 20 hybrids within a plot. Finally, we observe that mixtures have more yield stability compared to conventional plots, highlighting a new avenue for increased stability in higher risk environments. The lack of yield penalty and stability benefits are promising for future investigations of mixtures that may complement each other in disease resistance or abiotic stress tolerance and increase overall yield stability in the field.

## 1 INTRODUCTION

Monocultures have dominated annual cropping systems across commodity crops. This system has been driven by mechanization, improvement of crop varieties, and availability of synthetic fertilizer (Bullock, 1992; Power & Follett, 1987), allowing for increased production per hectare. However, such uniformity also has a drawback, as monocultures expose entire fields to significant risk of disease or pest outbreaks due to the lack of genetic diversity (Cook, 2006; McDonald, 2014; Stukenbrock & McDonald, 2008). More diversified cropping systems, such as intercropping where two or more crops are grown in the same field (Ofori & Stern, 1987), have long been proposed as an alternative solution to increase genetic diversity and reduce disease and pest pressure (Huss et al., 2022; Trenbath, 1993). However, intercropping has not been widely adopted in the US due to mechanical and management challenges (Bedoussac et al., 2015; Brooker et al., 2015; Glaze-Corcoran et al., 2020) and perceived farmer risk (Khanal et al., 2021; Rao & Singh, 1990).

Planting variety mixtures - mixtures of two or more varieties of the same crop species - offers a promising alternative by providing increased disease and pest resistance while maintaining a level of uniformity expected by the grower (Kopp et al., 2023; Wuest et al., 2021). For instance, Wuest (2021) demonstrated that wheat variety mixtures could reduce the incidence of disease. Similar effects have been observed in rice, where mixing varieties has improved resistance to certain pathogens (Zhu et al., 2000). The concept of refuge-in-the-bag where mixtures of seeds containing pest resistance genes and those without are sold directly to growers provides further precedent for the potential of variety mixtures (Yang et al., 2015).

While variety mixtures provide benefits for disease and pest resistance, in order to be implemented in a production field, mixtures must yield comparatively to a conventional system. In a large metadata study, Reiss & Drinkwater (2018) found that variety mixtures yielded 2.2% more than expected, and all crops evaluated except for sorghum experienced a significant yield increase. This benefit was more pronounced in environments with high disease pressure. It has been theorized that more functionally diverse variety mixtures occupy different ecological niches (Reiss & Drinkwater, 2018), and studies have shown that functional diversity contributes to the increase in mixture productivity over monocultures (Montazeaud et al., 2020; Su et al., 2024). The majority of the studies evaluated in the meta analysis were on small grains, and limited studies have been done on corn (Reiss & Drinkwater, 2018). A 1972 study showed that higher yielding maize lines within a variety mixture often contributes disproportionately more to yield than expected compared to a pure stand, and conversely lower yielding lines disproportionately less, but the overall variety mixture yield is not significantly different from the pure stand (Kannenberg & Hunter, 1972). This points to a potential avenue to improve disease resistance, reduce inputs (Schipanski et al., 2016; Tilman et al., 2011), and increase yield stability of maize through variety mixtures. However, the integration of variety mixtures within maize relies on the plants not outcompeting their neighbors, whether via shading from neighboring plants being taller or via nutrient capture of the roots.

It has been theorized that early selection pressure on individual plants within a breeding pipeline encourages competitive traits for resource capture, leading to intense competition among neighboring plants in later generations (Muir, 2005; Murphy et al., 2017). Conversely, it has been shown that the dramatic yield increases seen in modern maize have been partially driven by an increase in planting density and the resulting tolerance of modern hybrids to higher densities - which one could infer as plants being less impacted by neighbor competition (Duvick, 2005). Efforts have focused both directly and indirectly on the shade avoidance response, in which a plant grows taller in response to shading by a neighbor and may result in increased variability or reduced yield across a field (Jafari et al., 2024; Mansfield & Mumm, 2014). As such, plant breeders have long utilized two-row and four-row plots to limit neighbor interactions and competition from other genotypes due to the shade avoidance response, ideally producing field trials with less overall neighbor interference for yield (David et al., 2001).

Through the numerous density studies conducted, it is known that modern maize is tolerant to intra-genotype competition. This raises the question: How tolerant is modern maize to inter-genotype competition? Understanding these dynamics could have significant implications for how we conduct yield trials (e.g. 2- vs 4-row plots) and select and mix varieties in production fields. Here, we aim to address three primary questions: (1) Do neighboring genotypes impact the yield of focal plots, (2) How does this affect yield trials and selection practices, and (3) Is it possible to mix hybrids while maintaining or improving yield and yield stability?

To answer the above questions, we took two approaches. First, we leveraged historical data for both inbred and hybrid field trials to investigate the genetic effect of neighbors. Second, we conducted a three-year field experiment across multiple locations and a second experiment in three additional locations to evaluate the effects of mixing hybrids within the same plot on overall plot yield and yield stability both in two and four-row plots. We aim to provide a clearer understanding of neighbor competition and the potential for utilizing variety mixtures in modern agricultural systems.

## 2 MATERIALS AND METHODS

### 2.1 Between-plot interactions

#### 2.1.1 Historical data aggregation and preprocessing

Historical phenotypic and genotypic data from Genomes to Fields were used from the 2022 Prediction Competition dataset (Genomes To Fields, 2023), spanning years 2014 to 2021. These experiments were planted in two-row plots at standard agronomic planting densities, and extensive phenotyping was done for plant height and yield. Data were filtered to plots with neighboring plots that had genotypic data (e.g. not border rows) on both sides, and further filtered to field sites that had at least 300 plots that met this criteria, totaling 141 locations and 4381 genotypes.

Genotypic and phenotypic data for the NAM RILs were used from Hung et al., 2012. These RILs were previously grown as 2-row plots in 11 location-year environments in 2006 and 2007 and phenotyped extensively throughout each growing season, including plant height, leaf angle, and leaf area. Most fields were blocked by family, while one location-year was blocked by flowering time. Data were filtered to plots with neighboring plots that had genotypic data on both sides, and locations without trait measurements were removed. A total of 1,920 genotypes and 8 locations were included in the final analysis.

NAM RIL Hybrid genotype and phenotype dataset were used from (Ramstein et al., 2020), including the check genotypes. A subset of up to 80 RILs from each of the 24 NAM families were crossed onto a single tester PHZ51. The RILs that were crossed were selected by their flowering time, so that the latest of the earliest flowering families and the earliest of the latest flowering families were used. They were grown in 2-row plots at approximately 50,000 - 75,000 plants/ha in five locations across two years, with a total of eight environments.. Fields were blocked by NAM family, specifically to try to minimize light competition. Plant height, grain yield, and days to silk were measured across the locations. Data were filtered to plots with neighboring plots that had genotypic data on both sides, and further filtered to remove families with less than a maximum of 20 observations across all locations, resulting in 11 locations and 5,000 genotypes in single row plots for the final analysis.

#### 2.1.2 Genomic Data

In order to determine the genetic effects of neighboring plots, genomic data from each of the three historical datasets were processed for downstream analyses in a manner appropriate for each experimental design.

Genotypic data from Genomes to Fields (Genomes To Fields, 2023) was filtered for minimum and maximum allele frequencies of 0.005 and 0.995, and a site minimum count of 2250. A centered IBS genomic relationship matrix was calculated using Tassel (Bradbury et al., 2007).

NAM RIL genotypes were previously generated from Hung *et al* (Hung et al., 2012). To generate the genomic relationship matrix, sites were filtered to a minimum and maximum minor allele frequency of 0.01 and 0.99. Indels were removed, and 10 million random sites were selected to generate the kinship matrix.

In order to estimate neighbor effects without including the direct impact of neighbors, a leave-one-location-out approach was used with Echidna (Gilmour, A. R. (n.d.)) to calculate per-location BLUPs of each RIL in the following model:

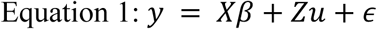

where y is an (N x 1) vector for all individual trait observations of a genotype. *X* is an (N x *r*) incidence matrix of *r* fixed effects for each observation. *β* is an (*r* x 1) vector of fixed effects for the environment. *Zu* represents the random effect associated with genotype where 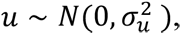 and *ε* is the residual error.

Hybrid genotypes were previously generated in Ramstein et al (Ramstein et al., 2020), except for the checks which were generated from hapmap3 using Tassel (Bradbury et al., 2007). The NAM RIL hybrids and check hybrid genotype tables were then merged by chromosome in Tassel (Bradbury et al., 2007). Each chromosome was filtered for a minimum and maximum minor allele frequency of 0.01 and 0.99, indels removed, and site minimum coverage of 950 genotypes. Chromosomes were then randomly sampled to 400,000 sites per chromosome and merged into one genotype table. A normalized IBS genomic relationship matrix was calculated using Tassel (Bradbury et al., 2007). GBLUPs were used from Ramstein et al., which were fit using a leave-one-location out method.

#### 2.1.3 Linear models for neighbor genotypic effects

Using lme4 (Bates et al., 2015) and lm packages in R (R Core Team, 2022), linear models were generated for each family and for each location in the NAM RILs and NAM hybrids. For plant height, the initial model was as follows:

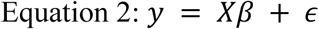

Where *y* is the (*N* x 1) vector of all individual observations of plant height. *X* is a (*N* x *r*) design matrix that contains *r* columns for the intercept, focal plot BLUP from equation 1, average neighbor plot BLUP from equation 1, and in the expanded model, additional trait BLUPs from equation 1. *β* is the (*r* x 1) vector of coefficients, and *ε* is the (*N* x 1) vector of residuals. This model was expanded to include neighbor BLUPs for leaf angle, leaf length, and leaf width. In the NAM hybrids, GBLUPs were used from Ramstein et al (Ramstein et al., 2020). The same model as described above was used for both height and yield.

#### 2.1.4 Mixed models for neighbor genotypic effects

The following model was fit using ASReml-R (Butler et al., 2023) for both plant height and yield across all experiments to estimate neighbor effects:

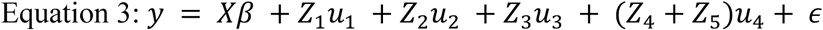

*y* is an (N x 1) vector of all individual observations of a genotype, such as plant height or yield.

*X* is an (N x *r*) incidence matrix of *r* fixed effects for each observation. *β* is an (*r* x 1) vector of fixed effects Field Location and Year. *Z*_1_*u*_1_represents the random genotypic effects of individuals, where 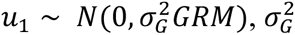 is the genetic variance and *GRM* is the genomic relationship matrix. *Z*_2_*u*_2_ represents the genotype-by-environment interactions of 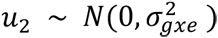 of each genotype and field location. *Z*_3_ is the design matrix representing field location and year combinations with 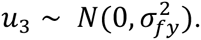 *Z*_4_is the design matrix representing the left neighboring genotype and *Z*_5_ is the design matrix representing the right neighboring genotype, where 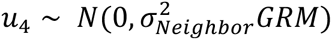 is the effect of having that genotype as a neighbor on either side.

The proportion of variance explained in the model by neighbors was done by extracting the variance components from the models: 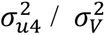 where V is the total variance explained in the model.

### 2.2 Within-plot interactions

#### 2.2.1 Genomes to Fields Two-Row Mixed Plot Experiment

A field trial was conducted in 2021 and 2022 to evaluate the yield of hybrids grown in mixture compared to those same hybrids grown as conventional single-hybrid plots within the larger Genomes to Fields experiment in 2020 and 2021. Twenty hybrid varieties from Genomes to Fields were selected for this experiment. These hybrids were developed from the same population and crossed on the same tester, had similar yield potential, and had varying heights and flowering time. Mixtures were designed as two full-factorial experiments, where each hybrid appeared in mixture ten times, totaling 50 plots. Fields were planted in 2021 in: Aurora, New York; Madison, Wisconsin; Lewiston, North Carolina; Champaign, Illinois; and Columbia, Missouri. In 2022, the same locations were included except for Illinois and Missouri. All locations except for Illinois had a single replicate, where Illinois had two. Each plot consisted of two rows and standard agronomic conditions were followed for each location as per the larger Genomes to Fields protocol (Genomes To Fields, 2023). In all locations except for North Carolina, plant and ear height were measured by taking a representative plant in approximately the 10th percentile, 50th percentile, and 90th percentile for each row. Days to anthesis and days to silking were recorded at the timepoint when 10%, 50%, and 90% of plants in a plot were flowering. Stand count, lodging count, and grain yield were measured at the end of the season for each plot according to the Genomes to Fields protocol (Genomes To Fields, 2023), with grain yield being mechanically harvested by a combine and moisture levels recorded.

#### 2.2.2 Four-Row Mixed Plot Experiment

In order to directly compare mixture yield with conventional plots in four-row plots, a second field trial was run in 2025. A different subset of 20 hybrids from Genomes to Fields with similar yield potential and as large of a height differential as possible were selected. Each hybrid appeared in mixture five times totalling 50 mixtures, as well as in a conventional single-hybrid plot. A “super mixture” plot consisting of 12 rows of all 20 mixtures was planted at the end of each replicate to evaluate the degree of competition when more than two varieties are planted. These plots, in addition to five check plots, were planted in two replicates at each location using a randomized complete block design. Fields were planted in St. Paul, Minnesota, Waseca, Minnesota, and Lamberton, Minnesota. Each plot consisted of four rows and standard agronomic conditions were followed for each location as per the larger Genomes to Fields protocol (Genomes To Fields, 2023). Plant and ear height were measured by taking a representative plant in approximately the 10th percentile, 50th percentile, and 90th percentile for each row. Days to anthesis and days to silking were recorded at the timepoint when 10%, 50%, and 90% of plants in a plot were flowering. Stand count, lodging count, and grain yield were measured at the end of the season for each plot according to the Genomes to Fields protocol (Genomes To Fields, 2023), with grain yield being mechanically harvested by a combine and moisture levels recorded.

#### 2.2.3 Data Analysis for Mixed Plots

Data was analyzed using R (R Core Team, 2022). Yield was calculated in bu/ac, adjusting for grain moisture at 15.5%, and then converted to MT/ha. Plant height GBLUPs for conventional, single hybrid plots within the G2F experiment were calculated using ASReml-R (Butler et al., 2023) using the following equation:

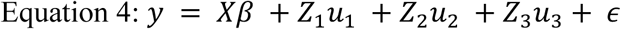

*y* is an (N x 1) vector of all individual observations of a genotype for plant height. *X* is an (N x *r*) incidence matrix of *r* fixed effects for each observation. *β* is an (*r* x 1) vector of fixed effects Field Location and Year. *Z*_1_*u*_1_represents the random genotypic effects of individuals, where 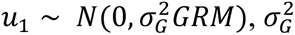 is the genetic variance and *GRM* is the genomic relationship matrix. *Z*_2_*u*_2_represents the genotype-by-environment interactions of 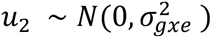 of each genotype and field location. *Z*_3_ is the design matrix representing field location and year combinations with 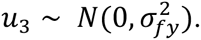

Within the 2025 four-row mixture experiment, yield BLUPs for conventional, single hybrid plots were calculated using ASReml-R (Butler et al., 2023) using the following equation:

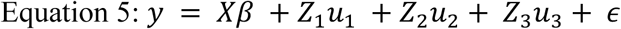

*y* is an (N x 1) vector of all individual observations of a genotype for yield. *X* is an (N x *r*) incidence matrix of *r* fixed effects for each observation. *β* is an (*r* x 1) vector of fixed effect Field Location. *Z*_1_*u*_1_represents the random genotypic effects, where *Z*_1_ is the incidence matrix linking observations to genotypes and 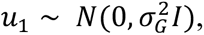 where 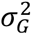 is the genetic variance. *Z*_2_*u*_2_represents the genotype-by-environment interactions of 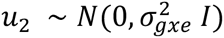 of each genotype and field location. *Z*_3_ is the design matrix representing replicate within each field location with 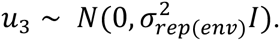

Predicted mixed plot yield was calculated as the average yield BLUPs of the two conventional single hybrid plots within that same location.

#### 2.2.4 Stability Analysis for Mixed Plots

A stability analysis of the 2021 and 2022 mixed plots was conducted using statgenGxE (van Rossum, 2025). Best linear unbiased estimators (BLUEs) were calculated in each location for hybrids both as a single hybrid plot and in each of the five mixtures. Outlier locations (North Carolina and Missouri) were excluded from the analysis. A Finlay-Wilkinson stability analysis was conducted, and the sensitivity recorded for each hybrid both as a single hybrid plot and within a mixture.

## 3 RESULTS

### Neighboring plot phenotypes vary widely across all experiments

In order to evaluate neighbor effects, multiple genotypes are needed that have a high degree of phenotypic variation (Figure 1). The maize nested association mapping (NAM) population consists of 25 families of 200 recombinant inbred lines (RILs) crossed onto a same recurrent parent, B73. The NAM RILs have a high degree of genetic variance, making them an excellent population to use for evaluating neighbor plot effects.The height differential between a focal plot and its two adjacent plots has a mean of 25 cm across locations, but can see a differential as great as 145 cm. When evaluating the height differential of a focal plot and its two neighbors, we find a correlation of 0.49. The NAM RILs are, however, limited by having a lower effective density compared to that of a hybrid production maize field due to reduced stature of the plants, and therefore reduced competition.

**Figure 1:**
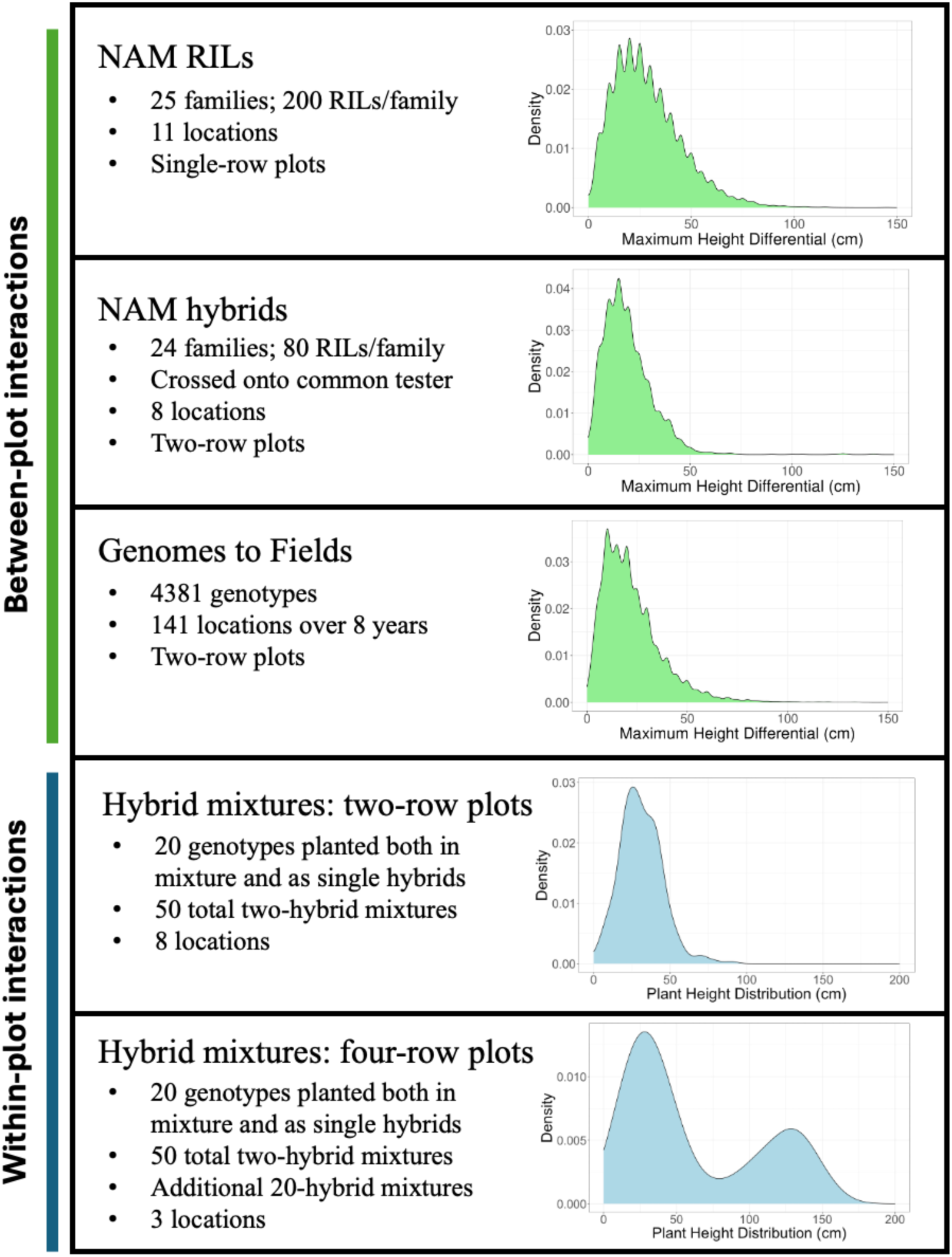
Overview of the five experiments analyzed and distribution of neighbor heights. In the across-row interaction experiments, the maximum height differential is the larger of the differentials between each neighboring plot and the focal plot. In the within-row interaction experiments, plant height distribution is the range of heights within a single mixed plot.

Crossing the NAM RIL lines to a common tester inbred allowed for the testing at agronomic density while still leveraging the genetic power of the NAM population. Across all families and locations, the mean yield of these plots was 6.56 tons/ha. The maximum height differential between neighboring plots ranges from 0 - 140 cm, with a median of 17 cm and a mean of 19 cm. The yield differential between neighboring plots ranged from 0.04 to 6.05 tons/ha, with a median and mean of 1.35 and 1.56 tons/ha, respectively. The total height differential between a focal plot and its neighbors and the yield of the focal plot had a correlation of 0.05.

Within Genomes to Fields, the average yield per plot across all years and locations was 9.84 tons/ha. Across all years, a range of 0 - 211 cm was observed with a median of 19 cm and mean of 22 cm between the heights of neighboring plots, while a range of 0 - 18.06 tons/ha in yield between neighboring plots was observed, with a median of 2.26 tons/ha and mean of 2.68 tons/ha. The total height differential and yield of the focal plot had a correlation of 0.01.

### Between-plot interactions

#### Neighbor height or yield has minimal impact on focal plot height or yield

In order to understand the degree in which neighboring plots impact focal plot height or yield, a simple linear model was implemented that predicts the focal plot trait based on the average traits of the neighbors. Across all of the environments in the NAM RILs, neighboring plot height accounts for an average of only 1.25% of the variance in focal plot plant height (t = 7.44, df = 7, p = 7.194e-05). Including more neighbor traits such as leaf angle, leaf width, and leaf length in the linear model increased the average variance explained to 2.6% (t = 11.051, df = 7, p-value = 5.515e-06). To evaluate the importance of neighbors compared to the focal plot in predicting focal plot height or yield, a β-ratio was calculated using the β for neighbor and focal plots within each model. The β-ratio indicates that there is family-based variation for neighbor importance (Figure 2); however, no statistically significant variation in neighbor importance is found when comparing heterotic groups or families (KW-test, p = 0.3246 and 0.4171). This lack of variation suggests that while there is substantial genetic variance in the NAM families, neighbor effects do not vary strongly across genetic backgrounds. When considering the coefficients (i.e. neighbor plant height, leaf angle, leaf width) in the linear models across NAM families and environments, the total number of significant effects surpass random expectation (binomial test, p = 0.0004). However, the specific traits impacting plant height vary widely by family, location, and year. These results indicate that environmental factors may have a greater influence on plant height than neighboring plants, further emphasizing the overall marginal effects from neighbors.

**Figure 2:**
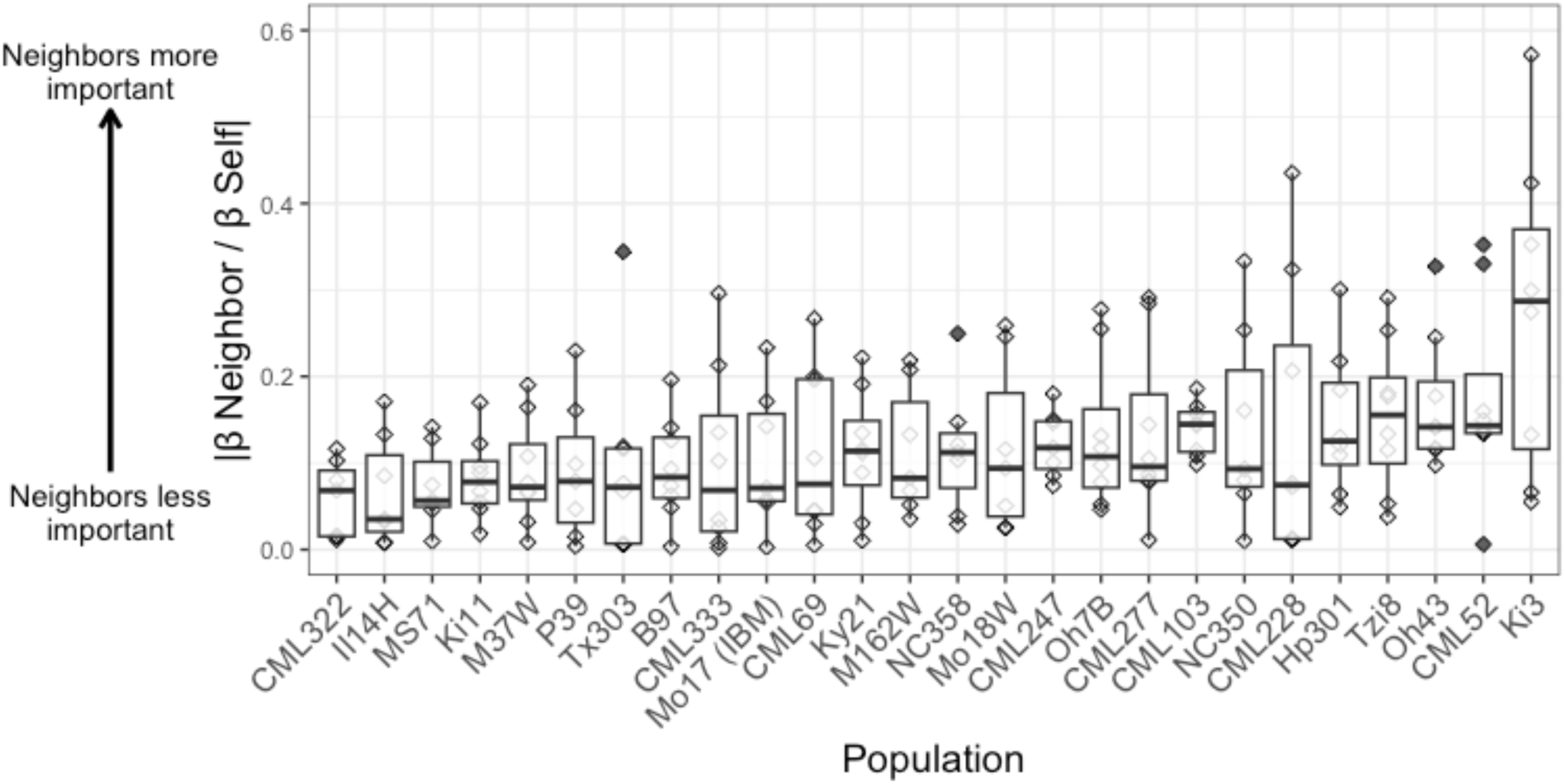
Distribution of neighbor β-ratios by family across all environments.

Yield is the ultimate measure of neighbor competition, and in order to measure it, hybrids are needed rather than inbreds. Building upon the same linear models as the NAM RILs, the NAM hybrids were utilized to predict focal plot yield. Yield of neighboring plots accounted for, on average, 1.7% of the variance in focal plot yield across environments (t = 10.232, df = 5, p-value = 7.655e-05). The combination of neighbor yield and neighbor height accounted for an average of 3.5% of the variance in focal plot yield across environments (t = 10.457, df = 5, p-value = 6.895e-05). The occurrence of significant effects for neighbor yield and plant height across all families and locations was slightly higher than random expectation, but was not significantly different (binomial test, p = 0.3173), again indicating limited neighbor effects.

#### Neighbor genetics explain 3% or less of the variance in focal plot yield

Linear models capture phenotypic neighbor effects on focal plot height or yield but are not able to account for the genetic contribution of neighboring plots. To address this limitation, we developed a mixed model that explicitly incorporates neighbor genetic effects through a genomic relationship matrix. In a single year (2021) of Genomes to Fields with 27 locations, neighbor genetic effects explained up to 3% of the variance in yield. However, across 141 location-year combinations environments, the genetic effects of neighboring plots account for only 1.55% of the explained variance in yield. This is a small but significant (SE = 0.007, Z = 8.78) effect. When predicting the height of a focal plot, neighbor genotype accounts for 0.19% of the explained variance (SE = 0.307, Z = 6.08).

The height of the neighbor plots was hypothesized to be contributing to the observed effect on yield of the focal plot based on the shading effects a taller neighbor may have on a shorter focal plot. To test the impact of neighbor height on the yield of a focal plot, we included the heights of neighboring plots into the model. This did improve the model fit (LR-statistic = 540.68, p = 0). Neighbor heights accounted for less of the explained variance than neighbor genotype, and were not statistically significant within the model (Z = 0.70).

Similarly, in the NAM hybrids, neighboring genotypes account for only 1.45% (SE = 0.007, Z = 3.31) of the variance in yield of the focal plot. Estimating different neighbor effects for each family resulted in a significantly better model fit compared to estimating the same neighbor effects across families (likelihood ratio test, p = 2.08e-05). This is likely due to the experiment being blocked by family in order to minimize light competition. Tzi8 was the only family to have significant neighbor effects (Bonferroni multiple test correction, Z > 2.87). Across all families, neighbor genotypes account for 0.5% of the explained variance in plant height (SE = 1.24, Z = 2.97). Incorporating family effects into the model identified the Tzi8 NAM family had significant neighbor effects (Bonferroni multiple test correction, Z > 2.87). The likelihood ratio test between the family and full neighbor models was not significant (p = 0.06), indicating only a marginal influence of family on the degree of neighbor effects on plant height. This again may be due to the blocking of the experiment by family.

In contrast, when evaluating the NAM RILs simultaneously across all families and environments for plant height using a competition mixed model, we find that neighboring genotypes account for 0.39% of the variance in plant height. Incorporating family effects into the model identified the Tzi8 and P39 families as having significant neighbor effects (Z > 2.87, Bonferroni correction), which we hypothesize could be due to these lines being selected under a lower planting density at their development. A likelihood ratio test between the two models with and without family effects was not significant (p = 0.83), again indicating that family does not have a major influence.

### Within-plot interactions

#### Mixture yield can be predicted based on conventional single hybrid plot yield

In a scenario with no competition effects, yield of a mixture would be the perfect average of the yields of the two hybrids in conventional, single hybrid plots. BLUPs were generated for each conventional hybrid in each location grown in 2025. Predicted yield was then calculated for each hybrid mixture in each location and had a correlation of 0.73 across all locations with actual mixed plot yield. Across all locations, the actual yield of the mixture was able to be predicted from the yield of the conventional plots with statistical significance (p = 5e-13, model p-val < 2.2e-16, adjusted r squared 0.55), indicating the potential for designing high yielding mixtures based off of single hybrid performance (Figure 3A).

**Figure 3:**
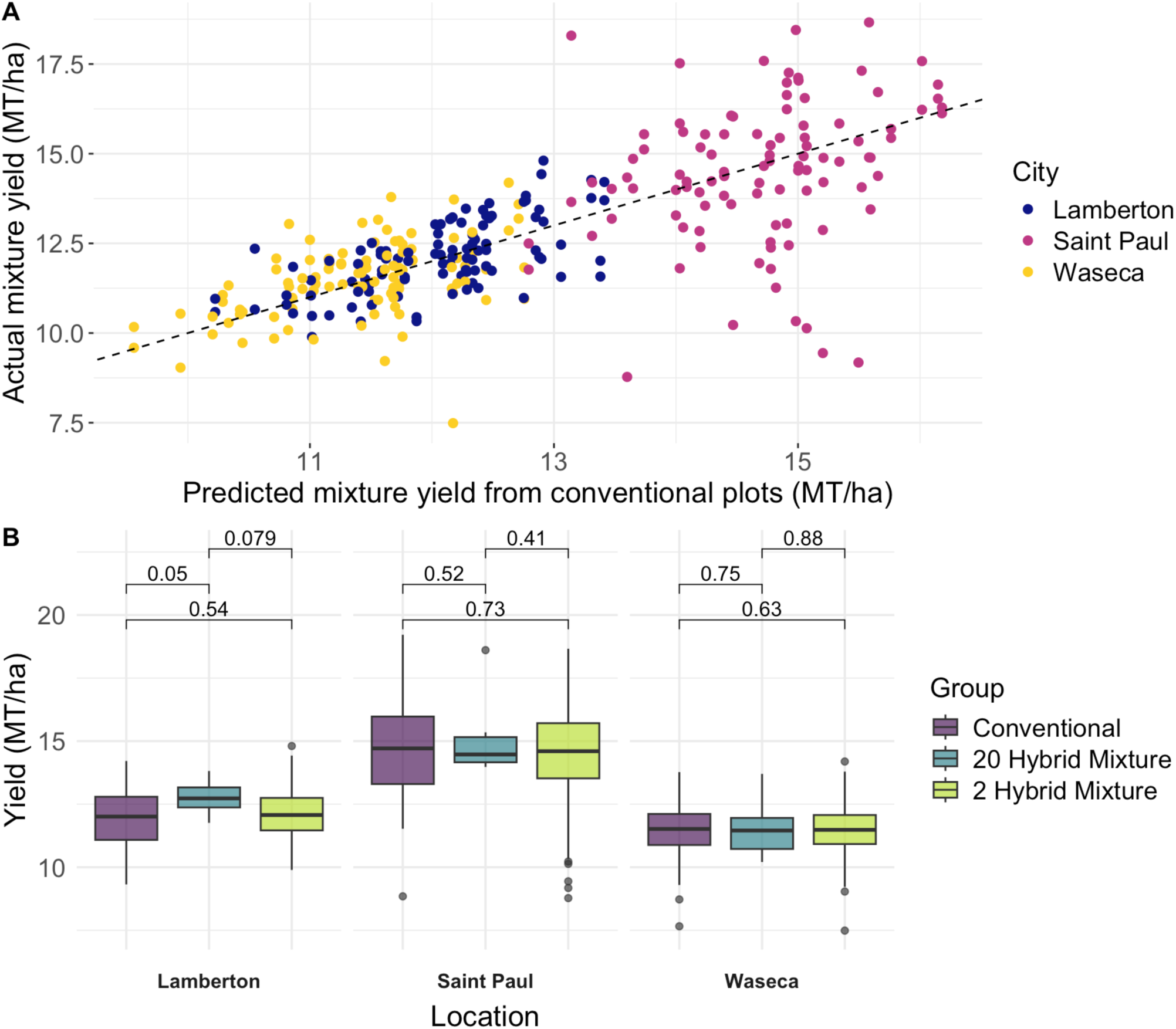
**A)** Predicted mixture yield from per-location BLUPs of the conventional plots compared to actual mixture yield. The dashed line indicates the 1:1 line where predicted yield equals actual yield. **B)** Mixtures of both 2 and 20 hybrids experience no yield penalty compared to the same hybrids grown in conventional single-hybrid plots.

#### Mixtures of up to 20 hybrids yield the same as conventional single-hybrid plots

Hybrid mixtures consisting of two hybrids provides a simple approach to understanding plant interactions within a plot. Increasing the number of hybrids into a “super mixture” of 20 hybrids results in greater phenotypic variation, potential differences in light and resource capture, and therefore greater neighbor competition. In 2025, three locations were planted with a 20-hybrid mixture and evaluated alongside conventional single hybrid and two hybrid mixtures. There were no significant differences in yield between the conventional, two hybrid mixture, and 20 hybrid mixture plots within each location, further highlighting the limited effects of neighbor competition in maize (Figure 3B).

#### Height differentials have no significant impact on mixed plot yield

Mixing hybrids with different heights but similar yield potentials provides another angle in which to evaluate neighbor effects in maize. Hybrid mixtures with large height differentials may experience more competition due to light capture than those that are more uniform in nature. We estimated the expected height differential of a mixed plot by taking the difference of the height GBLUPs for each hybrid grown as a single hybrid plot. This was done on a per-location basis.

The predicted height differential had a marginally negative effect on yield of the mixed plots across environments, although it was not significant (p-val = 0.06, model p-val < 2.2e-16). In the 2025 four-row plot experiment, the actual height differential (90th percentile - 10th percentile plant in plot) had no effect on yield across environments (p-val = 0.39, model p-val < 2.2e-16). This highlights that under modern agronomic conditions, height differentials have a limited impact on yield. On the other hand, some individual environments (North Carolina in 2021 and 2022; Wisconsin in 2021; St. Paul, MN in 2025) were significant in predicting yield, indicating that location effects have a larger impact on yield than the height differential of the mixtures.

#### Mixtures have increased yield stability compared to conventional single-hybrid plots

Yield stability is an important trait for growers, especially given the challenges of fluctuating environments. One hypothesized benefit of mixtures is yield stability. To evaluate this, we evaluated the yield variance in both conventional single hybrid plots and mixtures grown in the Genomes to Fields experiment in Wisconsin, New York, and Illinois. Two models were fit, the first allowing for homogeneous genotype-by-environment variance, and the second for heterogeneous genotype-by-environment variance between conventional and mixture (likelihood ratio test, χ² = 2.44, p = 0.12). We find a 49% reduction in genotype-by-environment variance in the mixtures (σ² = 201) compared to conventional plots (σ² = 293). This result indicates that across environments, the yield of mixtures is more similar (or, has an increased Type 1 stability), while conventional plots may perform well in one location and poorly in another, exhibiting Type 2 stability. A Finlay-Wilkinson stability analysis was conducted to directly evaluate this. While there was not a significant difference between the conventional and mixture plots across locations (p = 0.8242), a trend was observed where mixtures were more stable when they included the least stable hybrids (Figure 4). Further investigation into this response with additional experiments is warranted.

**Figure 4:**
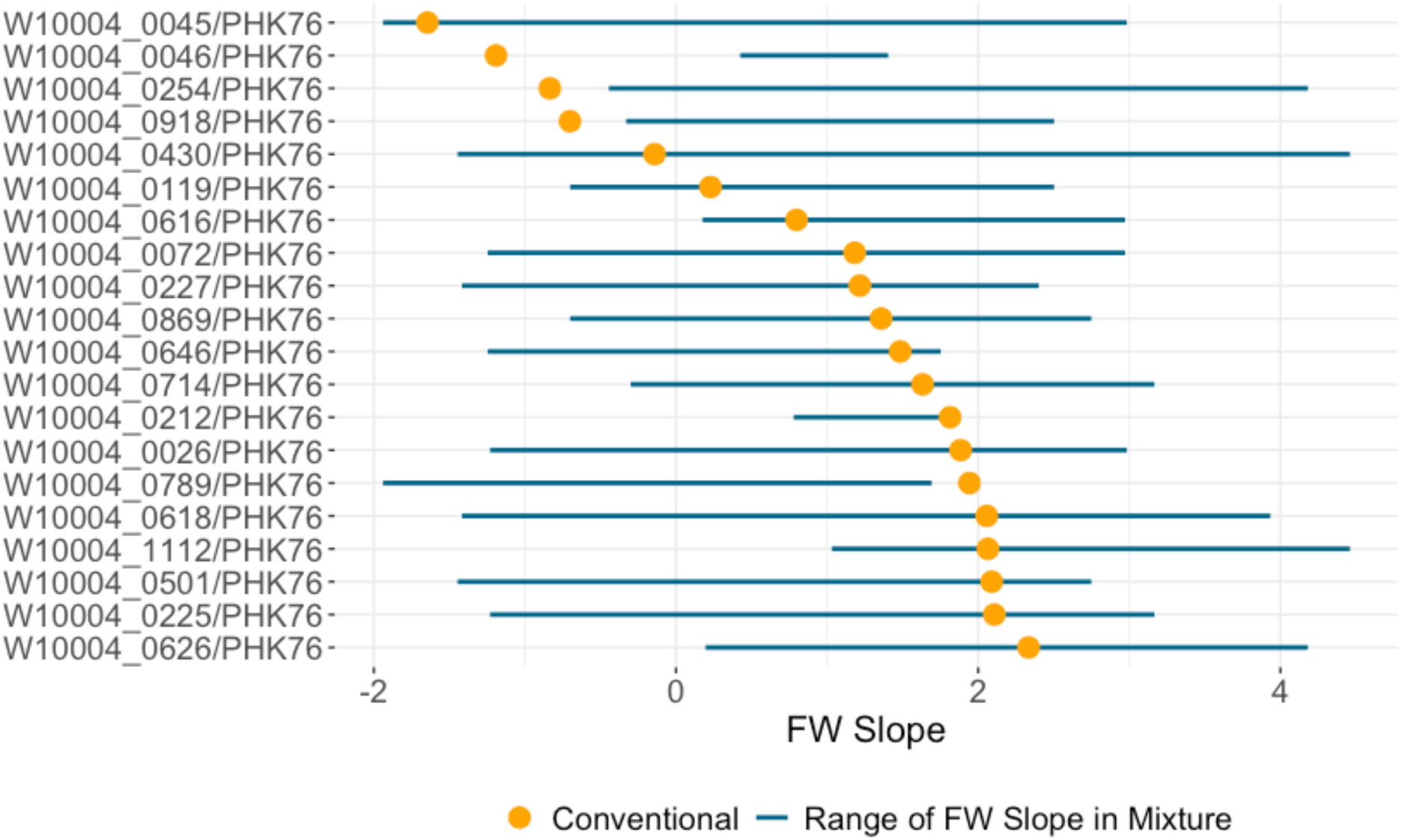
Stability metrics for each hybrid grown in a conventional single-hybrid plot and in mixture. A FW slope of 0 indicates perfect Type 1 stability, while values further away from zero have less Type 1 stability.

## 4 DISCUSSION

Overall, we find that neighboring plots resoundingly have marginal, but significant, effects on the focal plot’s height and yield. These trends occur across both single and two-row plots, and the genetic background of a variety plays a minimal role in neighbor effects.

Our findings across the NAM inbreds demonstrate that neighboring plots exert a small but significant influence on focal plots for plant height. Although these effects exceeded random expectation, the practical impact of neighbor traits is relatively minor and do not play a large role in determining plant height. This is further supported by the fact that the number of significant neighbor traits varied widely across family, location, and year, indicating that the magnitude of neighbor effects is context dependent. Notably, the 2007 New York location was blocked by flowering time and had nearly 3x fewer significant terms compared to the 2006 field site, which was blocked by family. While genotype-by-environment interactions certainly play a role in year-to-year variation across field sites, the other locations which were consistently blocked by family did not exhibit this magnitude of effect across years.

When extending these analyses to NAM hybrids, we found that within a family and location, the impact of neighboring plots on focal plots was not substantially greater than random expectation. Slightly less variation in the number of significant traits and locations compared to the NAM RILs suggests that modern hybrids, unlike inbreds, may be more resilient to competition or may have been bred to mitigate neighbor impacts, reflecting the yield gains achieved under increased planting densities. However, it is also important to note that the NAM hybrids have less genetic variation than the NAM RILs, and therefore could explain the reduction in variance seen.

In the Genomes to Fields (G2F) and NAM hybrid experiments, we observed marginal but significant neighbor effects on focal plots when predicting across years and environments, consistent for both yield and plant height in the competition mixed model. This underscores that neighbor genotype, though not a primary factor, does contribute to yield and plant height variance. Hybrid single-row plot trials in Ethiopia showed similar results, where the ratio of neighbor variance to the focal plot variance ranged from 4% to 10% across field trials (Keno et al., 2023). For comparison, our G2F hybrid experiment, using two-row plots, had a ratio of 3.7%. However, while competition effects can be modeled, they play a secondary role relative to genotype and environmental interactions in determining plant performance. Additionally, these results suggest that two-row plots are sufficient to minimize neighbor competition in early yield trials, which is contradictory to what a previous study found when measuring the impact of neighboring plot height differentials on the focal plot yield (David et al., 2001). The results from G2F suggest that neighboring plot plant height plays a smaller impact on the focal plot’s yield than previously reported.

Within the four-row mixed plot experiment, the relative predictability of mixed plot yield compared to the single hybrids was promising, and reflects previous studies (Kannenberg & Hunter, 1972). The hybrids within the mixed plot experiments exhibited similar yield potential but different plant architectures, providing unique insight into the effects of mixtures of varying heights on yield. Notably, the lack of yield response to varying height differentials in variety mixtures points to a promising strategy for improving yield stability, abiotic stress tolerance, and pest and disease resistance of the entire field through hybrid mixtures. For example, two potentially less stable hybrids could be mixed to develop a more stable mixture. Likewise, a high-yielding but more drought susceptible hybrid could be mixed with a drought tolerant hybrid with a slightly lower yield to improve stability without compromising yield. Successful mixtures need not be limited to only two hybrids and optimization of “super mixtures” should be investigated further for potential increases in yield stability.

Looking forward, incorporating these results into crop growth models could enhance our ability to predict yield responses to stressors when grown in mixtures. While neighbor effects remain an important factor to consider in certain populations and experimental designs, our results indicate that modern maize hybrids have been bred for increased tolerance to competition making them well suited to be successful in mixtures. Variety mixtures may offer a practical strategy for stabilizing yields and enhancing resilience. The potential for cooperative interactions among hybrids within variety mixtures offers promising avenues for breeding programs, paving the way for improved yield stability and resilience through variety mixtures.

## ACKNOWLEDGMENTS

This work was supported in part by the Minnesota Agricultural Experiment Station and the US Department of Agriculture–Agricultural Research Service (USDA-ARS) under Project Number 8062-21000-043-00-D (Plant, Soil and Nutrition Research Unit, Ithaca, NY), Project Number 5070-21220-046-000-D (Plant Genetics Research Unit, Columbia, MO), and Project Number 6070-21220-017-000-D (Plant Science Research Unit, Raleigh, NC). Mention of trade names or commercial products in this publication is solely for the purpose of providing specific information and does not imply recommendation or endorsement by the U.S. Department of Agriculture. USDA is an equal opportunity provider and employer. Part of this work was carried out by using the resources of the Cornell University BRC Bioinformatics Facility, which is partially funded by Microsoft Corporation. A.J.S. is supported by NSF GRFP DGE – 2139899 and NSF PRFB PGRP Award #2410349. We thank Guillaume Ramstein, Joe Gage, and Seren Villwock for their feedback on this manuscript.

## CONFLICT OF INTEREST

The authors declare no conflict of interest.

## DATA AVAILABILITY

All data and scripts will be made publicly available in our repository upon publication.

## Notes

### Competing Interest Statement

The authors have declared no competing interest.

